# Microstructural Impairments in a Topologically Distinct Prefrontal-Habenular Connection in Cocaine Addiction

**DOI:** 10.1101/2022.01.10.475656

**Authors:** Sarah G. King, Pierre-Olivier Gaudreault, Pias Malaker, Joo-won Kim, Nelly Alia-Klein, Junqian Xu, Rita Z. Goldstein

## Abstract

Drug addiction is characterized by neuroadaptations in mesocorticolimbic networks regulating reward and inhibitory control. The habenula (Hb) is central to adaptive reward and aversion-driven behaviors, serving as a hub connecting emotion/cognitive processing regions including the prefrontal cortex (PFC). However, its role in human drug addiction has not been fully explored. Using diffusion tractography, we detailed PFC structural connectivity with three regions, namely the Hb, ventral tegmental area (VTA), and anterior thalamus (AT), and quantified the tract-specific microstructural integrity using diffusion tensor imaging within the anterior limb of the internal capsule (ALIC) in healthy and cocaine-addicted individuals. White matter microstructure in cocaine-addicted individuals was uniquely impaired in PFC-Hb projections in the ALIC, distinguishable from adjacent PFC-VTA and PFC-AT projections, with more pronounced abnormalities in short-term abstinence. These findings extend preclinical evidence of PFC-Hb circuit impairments in addiction and contextualize the plausible existence of a similar PFC-Hb connection in the human brain.

## Introduction

Impaired reward valuation is a fundamental feature of drug addiction, which is characterized by compulsive drug-seeking at the expense of alternative reinforcers (*1*). Dysregulation of the prefrontal cortex (PFC), which is integral to the brain networks that regulate reward processing, salience attribution, and inhibitory control, is proposed to precipitate bingeing and relapse, perpetuating the addiction cycle (*2, 3*). The habenula (Hb), a reward-processing structure in the epithalamus, plays a critical role in linking emotion and cognitive processing regions, notably the basal ganglia/striatum and amygdala, but also the PFC (*4*–*6*). Within the reward system, the lateral nucleus of the Hb (LHb) conveys aversion-related information to monoamine-releasing nuclei, thereby regulating motivated behaviors and reward sensitivity (*7*). Specifically, neurons in the LHb trigger aversive responses to undesirable events, signaling to the rostromedial tegmental nucleus to suppress reward-sensitive neurotransmission in the dopaminergic midbrain (*8, 9*).

The LHb has been increasingly recognized as a key neural substrate of chronic pathological substance use (*10, 11*), with emerging preclinical evidence implicating it in drug-seeking and addiction. Specifically, certain LHb neurons show biphasic responses to cocaine, including an initial downregulation (marking a drug-induced “high”) followed by sustained increased firing (*12, 13*). Further, excitatory signaling from the rodent PFC to the LHb (*14*), and the LHb to the rostromedial tegmental nucleus (*15*), increased when cocaine-seeking behaviors were suppressed in a behavioral model of drug-related inhibition. Lasting LHb activation is consistent with aversive states that define withdrawal and, by promoting stress-induced cocaine-seeking, this mechanism may play a role in mediating the transition from initial use to long-term dependence, craving, and relapse (*16, 17*).

However, despite this unique role in cocaine addiction, few studies to date have investigated the specific PFC-Hb projection in the human brain, including in human drug addiction (*18*). A growing body of studies in small animal highlights unequivocal evidence for a monosynaptic medial PFC (mPFC) projection to the Hb (*19, 20*), which regulates socially directed behavior (*21*) and working memory (*22*). Although such a direct PFC-Hb projection has not been carefully validated in recent non-human primate tracer studies, a handful of studies in macaques do support the existence of a similar PFC-Hb anatomical connection in the primate brain (*4, 23, 24*). Additionally, the phylogenetically conserved nature of the Hb circuitry (*25*) further motivates the exploration of the PFC-Hb anatomical projection in humans with diffusion magnetic resonance imaging (MRI) tractography (*26*), taking advantage of the sizable afferent fibers to the Hb through the stria medullaris (SM), a relatively thick fiber bundle situated on the dorsal-medial surface of the thalamus (*6, 27*).

To evaluate the microstructural features of the PFC-Hb connection, it is preferred to restrict quantification to a common major fiber pathway to avoid partial volume effects. Although the exact anatomical pathway of the PFC-Hb in the human brain is still elusive, the anterior limb of the internal capsule (ALIC), a major fiber bundle that carries thalamic and brainstem tracts from the PFC, serves as a highly plausible conduit. White matter pathways with different functional origins are topologically organized within the ALIC (*28*). Furthermore, white matter pathology in the ALIC was previously observed in distinct PFC projections in humans with bipolar disorder, suggesting that microstructural hallmarks of diseases may be localized to specific tracts in this region (*28*). Similarly, the importance of white matter microstructural integrity to human drug addiction has been made evident through studies of diffusion tensor imaging (DTI), showing abnormalities in the ALIC (*29, 30*) as measured with reduced fractional anisotropy (FA) in major fiber tracts associated with the PFC (*31*).

The present study sought to delineate PFC-Hb structural connectivity through the ALIC and investigate its specific white matter microstructural features in individuals with cocaine use disorder (CUD), including short-term abstinent and current users, as compared to demographically matched healthy individuals. We first performed probabilistic diffusion MRI tractography using individualized Hb, and control region, seeds to assess the topology of structural connections with the PFC. Next, by probing the tissue microstructure using DTI, we aimed to elucidate the putative unique relationships between white matter microstructure in the PFC-Hb tract and drug addiction. Specifically, we hypothesized that PFC-Hb white matter microstructure is (1) characterized by reduced coherence in individuals with CUD as compared to healthy controls, and (2) associated with cocaine use severity measures, ascertained through both objective and self-report measures.

## Results

### Prefrontal-habenular tracts display distinct topology within and posterior to the ALIC

Qualitatively, the rending of structural connectivity prior to entering the ALIC showed the Hb streamlines follow a ventral-medial pathway relative to streamlines from the AT and VTA. Within the ALIC, the Hb streamlines maintain this ventral-/medial topology (Figure 2B). Such topological distinction was highly consistent across subjects. Quantitatively, the degree of overlap between binarized streamline tracts within the ALIC were less than 70%: for the Hb, 58.6% with AT and 52.9% with VTA; for the AT, 52.2% with Hb and 61.8% with VTA; and for the VTA, 52.8% with Hb and 69.8% with AT (Figure 2C).

**Figure 1.**
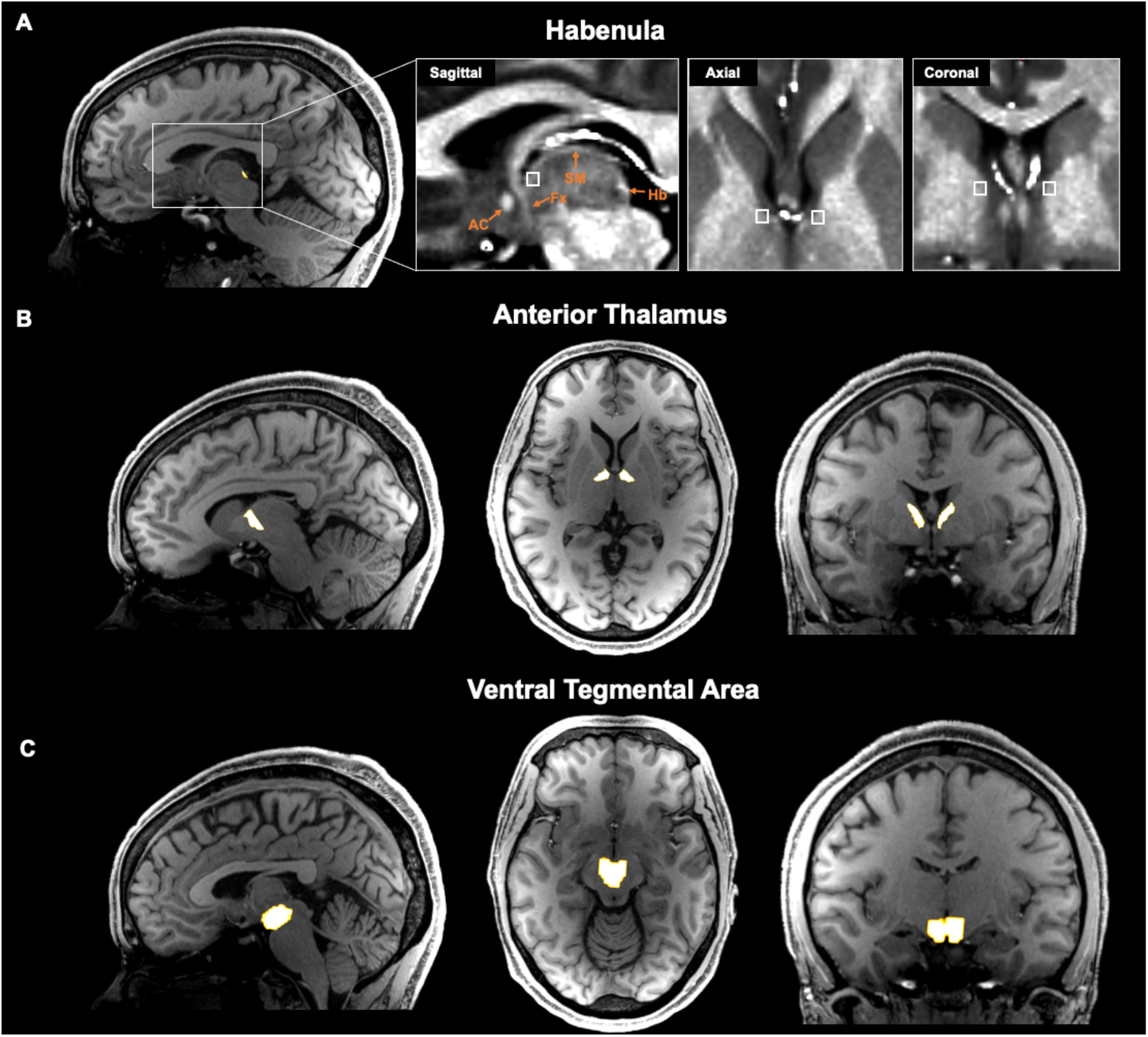
Representative subcortical seed region and inclusion mask selection for tractography analysis. (A) Habenula segmentation mask overlaid on T1w image. Anatomical landmarks are identified in the expanded boxed region with T1w/T2w image. Streamlines were constrained to pass through an individually defined anterior stria medullaris (white box) mask. AC=anterior commissure; Fx=fornix; Hb=habenula; SM=stria medullaris. (B) Anterior thalamus mask generated using Freesurfer’s anteroventral and ventral anterior nuclei segmentations. (C) Ventral tegmental area mask derived from an atlas of the midbrain in MNI space and warped to individual space.

**Figure 2.**
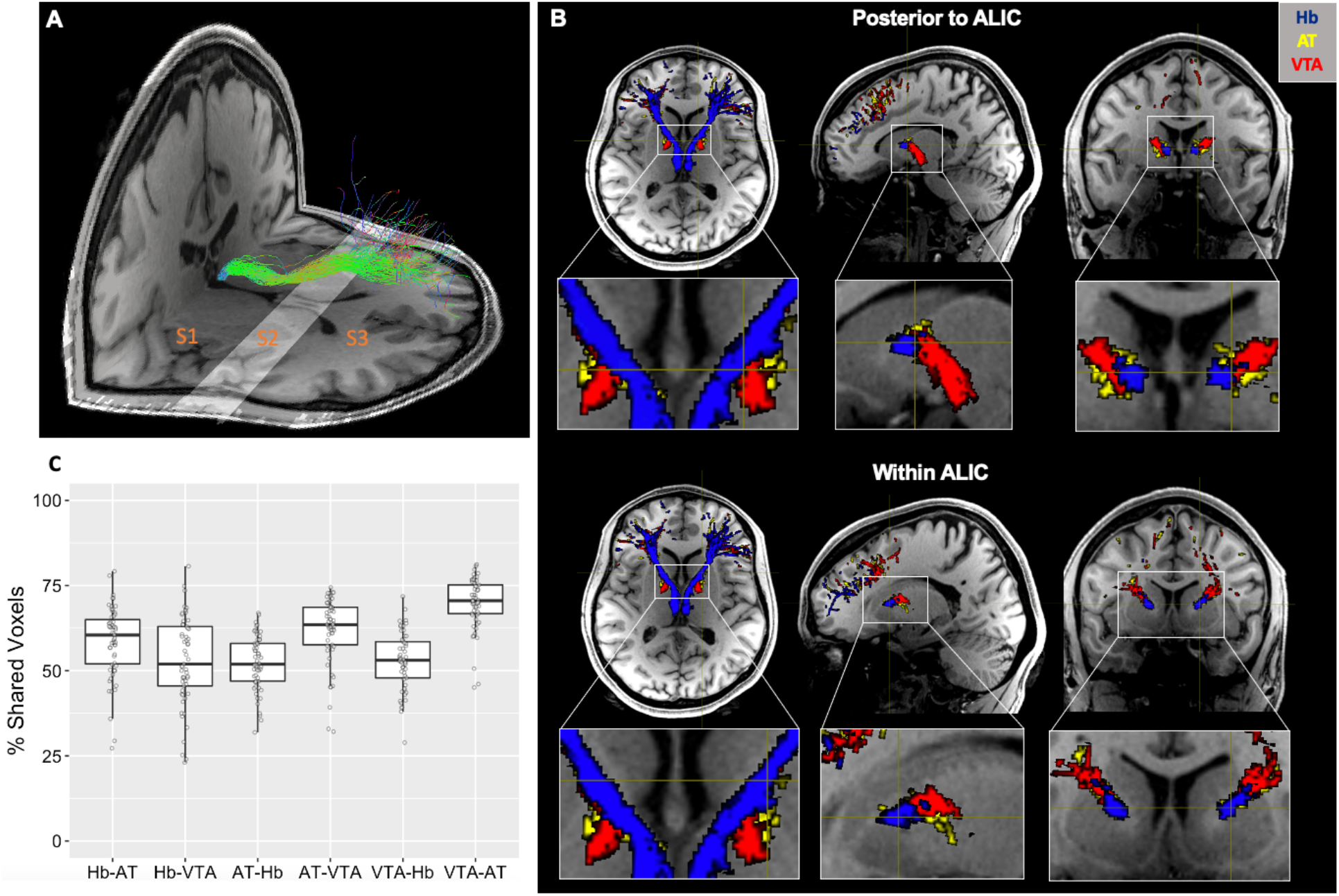
Representative habenula tract and its topological feature with the control tracts within and posterior to the anterior limb of the internal capsule (ALIC). (A) Designation of subsections using anatomical landmarks with habenula tracts (green) overlaid: S1, anterior of the habenula to anterior thalamus; S2, anterior of the thalamus to anterior caudate; S3, anterior of the caudate to PFC grey matter. (B) Topology of the habenula tracts (blue) immediately posterior to (upper panel) and within (lower panel) the ALIC (S2) compared to control AT (yellow) and VTA (red) tracts. The habenula tracts display a ventral-medial trajectory relative to the control tracts that is maintained throughout the ALIC. (C) Percentage of voxels shared between masks of each tract within S2 (e.g., Hb-AT denotes the percentage of Hb tract voxels that are also contained within the AT tract, see main text for detail). Boxplot represents the median (center line), interquartile range (box), and 1.5x the interquartile range (whiskers).

### Prefrontal-habenular tracts reach distinct prefrontal cortical target

All three subcortical seed regions produced streamlines terminating in the SFG, MFG, IFG, and OFC target regions in every scan, whereas fewer than two-thirds of scans produced any streamlines terminating in the FP or ACC (Figure 3B). As a result, statistical tests were performed only in the former four regions. The percentage of streamlines projecting from our seed regions of interest to the PFC target regions was analyzed using a 3 (Group: CTL, CUD+, CUD-) × 3 (Seed: Hb, AT, VTA) × 4 (Target: SFG, MFG, IFG, OFC) × 2 (Side: left, right) ANOVA. There was a significant main effect of Target (SFG > MFG > IFG > OFC), F(3,1344) = 556.45, p < .001, which was qualified by a significant Seed × Target interaction effect, F(6,1344) = 25.04, p < .001 (Figure 3B). Additionally, there was a significant Group × Target interaction effect, F(6,1344) = 2.74, p = .012 (Figure 3C). No other main or interaction effects reached significance, F < 1.14, p > .333. Post hoc analysis of the Seed × Target interaction showed a significant effect of Seed for all four targets and vice versa, implicating highly distinguishable targets in the PFC for the Hb streamlines as compared to the two control seed regions. For the effect of Seed: SFG (AT = VTA > Hb), F(2,348) = 40.84, p < .001; MFG (Hb > AT = VTA), F(2,348) = 7.02, p = .001; IFG (Hb > AT = VTA), F(2,348) = 3.51, p = .031; and OFC (Hb > AT = VTA) F(2,348) = 16.28, p < .001. For the effect of Target: Hb (MFG > SFG = IFG > OFC), F(3,464) = 158.41, p < .001; AT (SFG = MFG > IFG > OFC), F(3,464) = 244.81, p < .001; and VTA (SFG = MFG > IFG > OFC), F(2,464) = 211.23, p < .001. Finally, post hoc analysis of the Group × Target interaction showed a significant effect of Group in the MFG, the region containing the most Hb streamline terminals (CTL > CUD+ = CUD-), F(2,348) = 5.20, p = .006.

**Figure 3.**
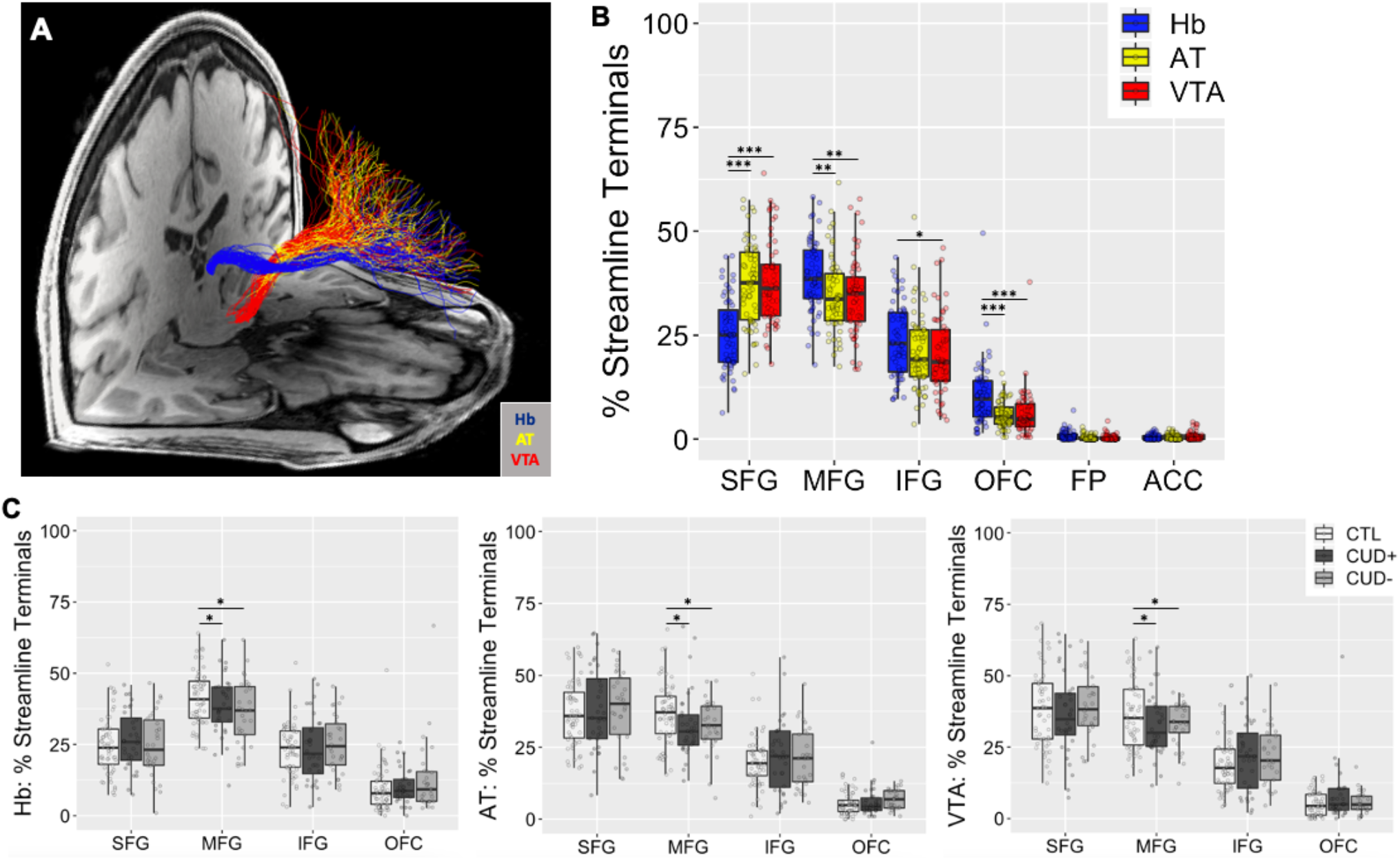
Distinct PFC-Hb streamline terminal distributions as compared to those from control seed regions. (A) Tracts seeded in subcortical seeds, the habenula (Hb, blue), anterior thalamus (AT, yellow), and ventral tegmental area (VTA, red), with streamline terminals in the PFC from a representative subject. (B) Comparison of the distribution of streamline terminals in the PFC for tracts seeded in subcortical seeds, averaged between left and right sides. *p < .05, **p < .01, ***p < .001. SFG = superior frontal gyrus; MFG = middle frontal gyrus; IFG = inferior frontal gyrus; OFC = orbitofrontal cortex; FP = frontal pole; ACC = anterior cingulate cortex. (C) CUD vs. CTL group comparisons of the distribution of streamline terminals from each of the subcortical seeds. Boxplots represent the median (center line), interquartile range (box), and 1.5x the interquartile range (whiskers).

### Microstructural integrity is impaired in PFC-Hb tracts in CUD compared to CTL

The four DTI metrics were compared separately, correcting for multiple comparisons, using 3 (Group) × 2 (Side) ANOVAs in the Hb tracts along the entirety of the streamlines terminating in the PFC, as well as in the AT and VTA tracts as controls (Figure 4A). For FA, this analysis in the Hb tracts revealed a significant main effect of Group (CTL > CUD+ = CUD-), F(2,112) = 6.06, p = .003, with no other significant main or interaction effects, F < 0.94, p > .333. This same analysis in the AT or VTA tract did not yield any significant effects after correcting for multiple comparisons, F < 4.77, p > .010. Additionally, there were no significant main or interaction effects for MD, F < 2.34, p > .101; AD, F < 1.66, p > .196; or RD, F < 3.69, p > .028, in any of the three tracts (Supplementary Figure 1A).

**Figure 4.**
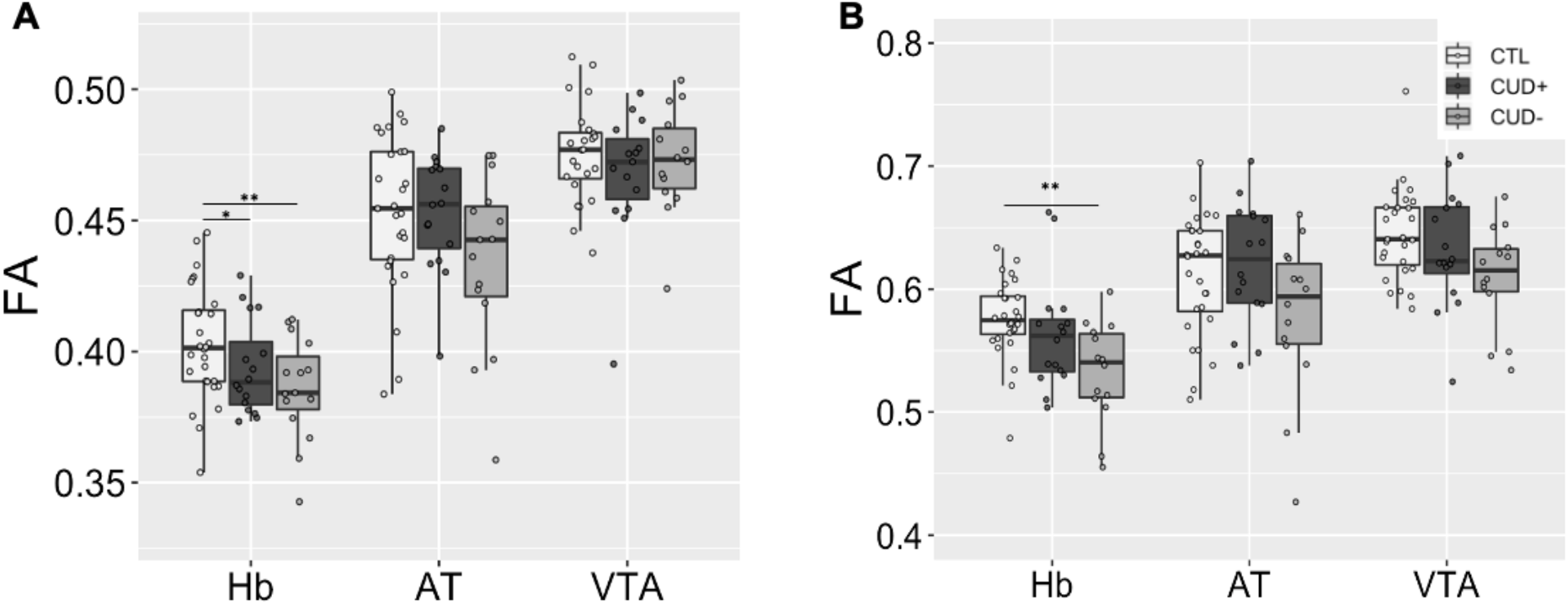
Group comparisons of DTI measured FA averaged across the entire tract (A) and within the ALIC subsection (B). *p < .05, **p < .01, ***p < .001. FA = fractional anisotropy; Hb = habenula; AT = anterior thalamus; VTA = ventral tegmental area; CTL = control individuals; CUD+ = currently using individuals; CUD-= short-term abstinent individuals. Boxplots represent the median (center line), interquartile range (box), and 1.5x the interquartile range (whiskers).

Additional analyses were performed separately in the ALIC (S2) subsection of each of the tracts to assess the specificity of the DTI metrics in a majority white matter region where the three pathways converge using 3 (Group) × 2 (Side) ANOVAs (Figure 4B). For FA, this analysis in the Hb tract revealed a significant main effect of Group (CTL > CUD-), F(2,212) = 6.66, p = .002, with no other significant main or interaction effects, F < 1.82, p > .166. This same analysis in the AT and VTA tract did not yield any significant results after correcting for multiple comparisons, F < 4.62, p > .012. Additionally, there were no significant main or interaction effects for MD, F < 1.16, p > .318; AD, F < 1.44, p > .242; or RD, F < 4.70, p > .011 (uncorrected trend), in any of the three tracts (Supplementary Figure 1B).

Finally, to assess the segmental specificity of the DTI group effect for FA in the Hb tract, we performed three 3 (Group) × 2 (Side) ANOVAs on FA in S1 and S3 in addition to S2 (see above), with Bonferroni correction for the three separate comparisons (alpha level .05/3 = .017) (Figure 5). This analysis yielded no significant main or interaction effects in either S1 (F < 0.73, p > .484) or S3 (F < 1.92, p > .150).

**Figure 5.**
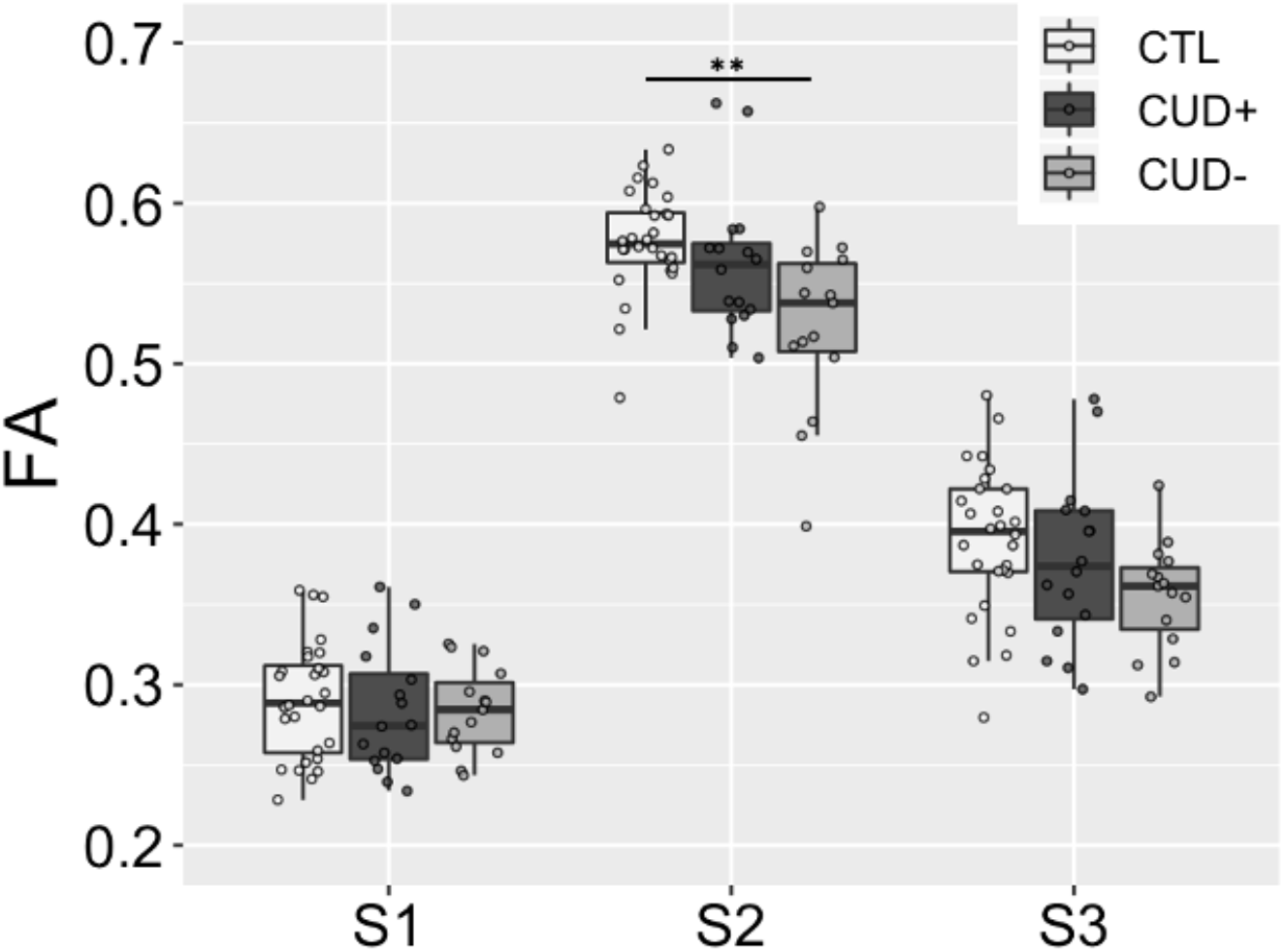
Group comparison of DTI measured FA in subsections S1, S2, and S3 of the PFC-Hb tract. FA = fractional anisotropy; CTL = control individuals; CUD+ = currently using individuals; CUD-= short-term abstinent individuals. Boxplot represents the median (center line), interquartile range (box), and 1.5x the interquartile range (whiskers). Note that S2 in this figure and Figure 4B Hb show the same data.

To inspect for potential covariates, correlations were performed between the DTI measures showing significant group effects (FA in the entire Hb tract and in the ALIC specifically) and demographic variables that differed significantly between the groups (i.e., years of education, verbal and non-verbal intelligence, depression symptomatology, nicotine dependence, and alcohol use); we also inspected group differences in these microstructural measures using “cigarette use” as a categorical variable. Additionally, we performed correlation analyses with the percentage of streamline terminals in the MFG, since this measure also differed significantly between groups. None of these tests were significant and, therefore, these variables were not included as covariates in the main analyses.

### Microstructural integrity in PFC-Hb tracts is not correlated with self-reported measures of cocaine use history and addiction severity

Two factors were extracted from the exploratory factor analysis (Factor 1 SS loading = 2.57, Factor 2 SS loading = 1.53) accounting for 28.6% and 17.0% of the total variance, respectively. Factor 1 loadings weighed most heavily on measures of recent cocaine use and current addiction symptomatology, specifically “duration of current abstinence” (factor loading = 0.988), “use in the past 30 days” (factor loading = -0.636), severity of dependence (factor loading = 0.664), craving (factor loading = -0.595), and withdrawal (factor loading = -0.459), with only weak contributions from other variables (factor loading < |0.39|). Factor 2 was conversely associated with more chronic drug use measures, with the highest loadings from “duration of heaviest use” (factor loading = 0.997) and “duration of regular use” (factor loading = 0.532), as well as “duration of longest abstinence” (factor loading = 0.405). Extracted scores from Factor 1 were sign-inverted to reflect a “recency” dimension (i.e., higher scores correspond to more recent drug use), whereas the raw scores from Factor 2 were considered to reflect a “chronicity” dimension (i.e., higher scores correspond to a more chronic drug use history).

Linear regression analysis was performed in the entire CUD group to assess correlations between these drug use factors and mean FA for both the entire PFC-Hb tract or within the ALIC, which was averaged between the left and right hemisphere due to the absence of any lateralization effects. This analysis did not yield any significant correlation with either recency, p = .614, or chronicity, p = .287, of cocaine use.

## Discussion

For the first time in the human brain, using anatomically constrained probabilistic diffusion tractography, we present consistent evidence for a plausible PFC-Hb connection. Although the existence of the PFC-Hb tract in the primate brain is not well established, a growing body of small animal evidence points to an unequivocal monosynaptic medial PFC-LHb projection (*19*–*21, 32*), which is consistent with the preferential MFG termination of our Hb streamline results (*22*). In addition, the ventromedial topological feature of our PFC-Hb tract is consistent with the well-known, phylogenetically conserved midline orientation of the Hb circuit (*6, 33*). Furthermore, tracer studies in macaques with relatively broad cortical coverage of the anterograde tracer injection sites, specifically from (i) Brodmann area 32/25/24b (*4*), (ii) orbitofrontal cortex (*24*), and (iii) Brodmann area 9/10 (*23*), show evidence of direct anatomical connectivity to the Hb. This converging evidence prompted us to adopt the term “PFC-Hb tract” to describe our current noninvasive neuroimaging results; a strict neuroanatomical validation would require careful replication of the tracer studies in macaques to validate whether the PFC-Hb pathway is conserved in the human brain. However, with limited tracer studies available in nonhuman primates to date, we recognize the on-going debates as to the existence, as well as the precise anatomical location, of the intended fiber pathways analyzed in this study.

Nevertheless, motivated by the specificity of the PFC-Hb projection, we quantitatively evaluated the microstructure of these pathway in cocaine-addicted individuals grouped by recency of use, as measured objectively by urine toxicology on the study day (dividing current vs. short-term abstinent users). In support of our first hypothesis, the PFC-Hb tract displayed decreased FA in both of the addicted subgroups compared with healthy controls, driven by the ALIC sub-section (S2). Similarly, using voxel-wise whole-brain analysis methods, white matter abnormalities, mostly in the form of diffuse reductions in FA, encompassing major cortical tracts (superior longitudinal fasciculus, corpus callosum, and anterior thalamic radiations) were previously reported in cocaine addiction (*30, 34, 35*) (although some studies have also reported increases in FA) (*36*–*38*). Consistent with our second hypothesis, the PFC-Hb FA was associated with an objective measure of recent cocaine use (urine toxicology), interestingly with abstinent individuals (abstinence < 6 months), displaying the most pronounced reductions as compared with healthy controls, while the current users displayed an intermediate phenotype. These subgroup differences may reflect neuroadaptive processes during abstinence, consistent with a previous report of decreased FA in the ALIC in short-term (< 1 year) but not long-term (> 1 year) abstinent addicted individuals (*30*). In general, this interpretation is also consistent with the progressive increase in cue-induced drug craving, or incubation of craving, as well as synaptic plasticity mechanisms that show a non-linear effect such that they peak in the early to mid-stages of abstinence and dissipate over longer time scales (*39*). Unexpectedly, DTI measures of white matter abnormalities did not correlate with any of the other, self-reported, measures of cocaine use history and addiction severity, including measures of both recent and lifetime use, in our exploratory analysis. Overall, whereas previous studies in drug addiction have primarily focused on large white matter fiber bundles, our hypothesis-driven, topologically differentiated results within the ALIC allowed us to observe significant differences in DTI metrics specific to the PFC-Hb tract that were not seen for other proximal seed regions, even after stringent statistical testing. Our results therefore enhance the specificity of circuit-based analysis for small tracts of particular importance to drug addiction, with potentially greater sensitivity to specific addiction sub-phenotypes.

The link between cocaine use and abnormalities in white matter FA is consistent with cocaine-induced neurotoxicity and neuroinflammatory effects, including reactive gliosis, which may disrupt the microarchitectural organization of axon bundles especially in subcortical regions (*31, 40*). Additional mechanisms underlying these white matter alterations in cocaine addiction may encompass elevated risk for cerebrovascular events, including ischemic and hemorrhagic stroke, as have been previously reported in frequent cocaine users, notably in the cerebral arteries that supply the subcortical white matter (*41*–*43*). Increased vasoconstriction can also potentially impact the myelination and structural integrity of axons in the affected fiber bundles (*44*). Although the direct measurement of these physiological processes on a microscopic scale is not possible with DTI, in a rodent model of cocaine addiction reduced white matter FA and increased RD were specifically colocalized with myelin damage and destabilized neurite outgrowth in the internal capsule and the corpus callosum (*45*). Our results in the CUD groups showing significantly decreased FA in the ALIC of PFC-Hb tracts, in combination with a trend towards increased RD (p = .011 uncorrected), may similarly reflect greater axonal dispersion or lower packing density resulting in less coherent directionality of the diffusion along these pathways.

Our streamline termination results suggest that, compared with the other subcortical regions examined, the Hb exhibits stronger connectivity with the cortical regions that are associated with decision-making and inhibitory control (i.e., the dorsolateral and ventrolateral PFC), which supports further interrogation of the functional significance of Hb connectivity in addiction. As an incidental finding (not correlated with our main results), we observed reduced streamline terminal density in the MFG relative to the other PFC subregions for all three subcortical seed regions of interest in both CUD groups. The dorsolateral PFC, which is located within the MFG, serves important roles in learning, memory, and higher order executive functions such as cognitive control/shifting, and is frequently reported in relation to impulsivity and motivated behaviors in drug addiction (*3*). However, the anatomical connectivity profile of the dorsolateral PFC in humans has yet to be fully characterized. Effective measurement of its structural connection strength might be attainable using advanced quantitative methods such as the apparent fiber density (*46*), which are beyond the scope of the current study. Further investigation of the possible role of dorsolateral PFC limbic connectivity in drug addiction is nevertheless warranted.

In contrast to Safadi et al., 2018, which also targeted the ALIC in humans, we performed our tractography from separate subcortical seed regions to cortical targets. The tissue composition of these subcortical seeds is complex, as are the axonal projections/terminals associated with these structures. This is further complicated by the limited image resolution of in vivo diffusion tractography, especially where different tracts organize as they leave the ALIC to reach a mixture of targets. As a result, the three tracts in this study may inevitably be contaminated by adjacent structures that project to/from the PFC through the ALIC. Specifically, our neuroimaging-defined PFC-Hb tract probably represents neuroanatomical projections in the dorsal-posterior direction at the level of the thalamus, which may contain the medial mediodorsal thalamic nucleus, although to the best of our knowledge there is no evidence that this region receives projections through the SM (*27, 47*). Furthermore, our PFC-VTA tract likely represents multiple ascending neuroanatomical projections from the inferior direction in the brainstem, and the PFC-AT tract may also represent projections from other thalamic nuclei passing through the AT. Recognizing this inevitable gap between in vivo diffusion tractography and the underlying neuroanatomy, the strong topological consistency of our results nevertheless adds an important piece of new information to the field, urging further tracer studies in non-human primates of the PFC-Hb tract.

The imaging acquisition methods and resolution employed here also prohibited separate segmentation of the lateral and medial Hb, both of which display notable functional distinctions in the context of addiction (*10*). Recent ultra-high resolution multi-shell diffusion MRI acquisition methods may allow for greater precision in tract estimation by reducing the potential for contamination from medial Hb fibers (*48*). Nevertheless, PFC input to the Hb terminates exclusively in the lateral nucleus (*49*), which supports our decision to emphasize functions attributed to the LHb. Additionally, a more complete interrogation of the Hb’s circuitry, including of its connections to the dopaminergic and serotonergic midbrain, will be vital for understanding the downstream effects of impaired PFC connectivity on the structure and functioning of the larger reward system. Finally, future studies with larger sample sizes and a longitudinal design, and objective measures of drug use, may better discriminate the effects of abstinence and recovery on the role of the PFC-Hb projection in drug-seeking and other drug-biased behaviors.

In summary, using in vivo diffusion tractography we found a topologically distinct fiber pathway between the PFC and Hb in the human brain that parallels the PFC-Hb projections reported in small animal studies. Using DTI measures obtained specifically along this PFC-Hb projection, we show microstructural impairments in individuals with CUD, as well as more severe abnormalities in this tract within the ALIC in abstinent users, which may reflect early neuroadaptations during short- to mid-term abstinence. Overall, our results advance ongoing research in the field by targeting a previously unexplored circuit in the pathophysiology of cocaine addiction in humans. Importantly, the consistency between the preclinical literature and our results supports further interrogation of the PFC-Hb tract as a neural target for better understanding the impact of individual differences in drug use and abstinence on reward processing, salience attribution, and inhibitory control in drug addiction.

## Materials and Methods

### Participants

Thirty-seven individuals with CUD and thirty-seven healthy control (CTL) subjects were recruited for the current study. Cocaine-addicted individuals were considered for inclusion if they had a history of CUD as assessed with diagnostic interviews that included the Structured Clinical Interview for the *Diagnostic and Statistical Manual of Mental Disorders, Fourth Edition* (*50*) and the Addiction Severity Index (*51*). Symptoms of cocaine withdrawal, craving, and severity of dependence were evaluated with the Cocaine Selective Severity Assessment (*52*), the 5-item Cocaine Craving Questionnaire (*53*), and Severity of Dependence Scale (*54*), respectively. Cigarette use and nicotine dependence were assessed with the Fagerström Test for Nicotine Dependence (*55*). A urine toxicology test completed by all participants was used as an objective assessment of drug use recency, which was further used to classify the CUD group as current users/cocaine urine-positive (CUD+) or abstinent users/cocaine urine-negative (CUD-). The mean current abstinence duration on the day of the MRI scan was approximately three days for the CUD+ group and three months for the CUD-group. Verbal and non-verbal intelligence were estimated using the reading subtest of the Wide Range Achievement Test 3 (WRAT-3) (*56*) and the Matrix Reasoning subtest of the Wechsler Abbreviated Scale of Intelligence (WASI) (*57*), respectively.

All individuals with CUD met criteria for cocaine dependence. Other comorbidities included alcohol use disorder (n = 13), cannabis use disorder (n = 5), opiate use disorder (n = 5), polysubstance use disorder (n = 2), intermittent explosive disorder (n = 1), and post-traumatic stress disorder (n = 1). All substance use disorder comorbidities were in partial or sustained remission at the time of the study. The CTL participants did not meet criteria for any of these disorders. Across all subjects, exclusion criteria were (1) history of head trauma or loss of consciousness (30 minutes or longer), or any neurological disease; (2) abnormal vital signs at the time of screening; (3) history of major psychiatric disorders, including current abuse/dependence for any substance with the exception of cocaine for the CUD group, disorders of high comorbidity with CUD (e.g., post-traumatic stress disorder, polysubstance use disorder), and nicotine/caffeine for all participants; (4) severe levels of self-reported depression as assessed using the Beck Depression Inventory (*58*) (scores > 20); (5) positive breathalyzer test for alcohol or positive urine screen for any psychoactive drugs, with the exception of cocaine in CUD participants; (6) current use of any medication that may affect neurological functions; and (7) MRI contraindications including any metallic implants, pacemaker device, or pregnancy.

Further decisions on participant exclusions were made based on MRI quality assurance. Scans were excluded based on excessive motion artifact in diffusion-weighted images (*n* = 1 CUD; *n* = 1 CTL); signal dropout in anterior cortical regions (*n* = 2 CUD); and incidental findings as indicated by a radiologist (*n* = 1 CTL). Scans from which tractography streamlines were successfully generated in the primary tract of interest (endpoints in Hb and PFC) were included in further analyses for a final sample of *n* = 28 CTL and *n* = 31 CUD (16 CUD+, 15 CUD-) (i.e., insufficient streamlines were generated for *n* = 7 CTL and *n* = 3 CUD). Participant characteristics are summarized in Table 1.

**Table 1.** Demographic, psychometric, and substance use characteristics of study participants^a^

|  | CTL<br>N = 28 | CUD+<br>N = 16 | CUD-<br>N=15 | Test Statistic |
| --- | --- | --- | --- | --- |
| **<u>Demographics</u>** |  |  |  |  |
| Age | 41.7 ± 7.5 | 46.8 ± 7.5 | 45.5 ± 9.4 | F = 1.86 |
| Sex (female/male) | 10/18 | 4/12 | 4/11 | $\chi^2 = 1.23$ |
| Race (Black/Other<sup>b</sup>/no response) | 22/4/2 | 14/1/1 | 8/6/1 | $\chi^2 = 6.52$ |
| Ethnicity (Hispanic/Non-Hispanic/no response) | 4/23/1 | 2/12/2 | 3/11/1 | $\chi^2 = 0.35$ |
| Education (years) | 14.3 ± 2.1<sup>2,3</sup> | 12.4 ± 2.1<sup>1</sup> | 11.9 ± 2.0<sup>1</sup> | F = 8.67* |
| **<u>Psychometrics</u>** |  |  |  |  |
| WRAT-3 Reading Scaled Score | 100.8 ± 10.8<sup>2</sup> | 89.6 ± 11.9<sup>1</sup> | 95.9 ± 13.3 | H = 8.72* |
| WASI Matrix Reasoning Scaled Score | 10.6 ± 2.8<sup>2</sup> | 8.3 ± 3.4<sup>1,3</sup> | 11.1 ± 2.8<sup>2</sup> | H = 7.75* |
| Beck Depression Inventory | 2.3 ± 3.0<sup>2,3</sup> | 7.3 ± 7.0<sup>1</sup> | 8.1 ± 8.4<sup>1</sup> | H = 12.20* |
| **<u>Substance use</u>** |  |  |  |  |
| Past 30-day alcohol use (# of days) | 1.2 ± 3.9<sup>2</sup> | 4.8 ± 3.2<sup>1</sup> | 1.0 ± 3.5<sup>2</sup> | H = 10.85* |
| Cigarette use (current/past/never) | 2/6/20 | 15/1/0 | 11/3/1 | $\chi^2 = 38.69^*$ |
| Fagerström Test for Nicotine Dependence | 0.8 ± 1.6<sup>3</sup> | 1.5 ± 2.4 | 2.9 ± 2.2<sup>1</sup> | H = 11.07* |
| Age of first cocaine use (years) | --- | 20.2 ± 5.4 | 23.3 ± 4.9 | t = 1.65 |
| Duration of regular cocaine use (years) | --- | 23.4 ± 10.0 | 16.5 ± 8.7 | U = 71.5 |
| Duration of heaviest cocaine use (years) | --- | 13.4 ± 12.7 | 11.1 ± 10.6 | U = 109.5 |
| Duration of longest abstinence (years) | --- | 4.8 ± 5.3 | 4.9 ± 6.8 | U = 96.0 |
| Duration of current abstinence (days) | --- | 3.3 ± 3.5 | 89.2 ± 62.1 | U = 24.5* |
| Past 30-day cocaine use (# of days) | --- | 13.4 ± 8.4 | 3.4 ± 7.0 | U = 28.5* |
| Cocaine Selective Severity Assessment | --- | 21.8 ± 4.1 | 16.3 ± 12.5 | U = 91.5 |
| Cocaine Craving Questionnaire | --- | 21.6 ± 13.1 | 8.3 ± 11.5 | U = 50.5* |
| Cocaine Severity of Dependence Scale | --- | 4.3 ± 3.6 | 8.2 ± 5.8 | U = 169.5 |

<sup>a</sup> $\chi^2$ tests were used for categorical variables; ANOVA or Student's t-test for normally distributed continuous variables; and Kruskal-Wallis H or Mann-Whitney U for non-normally distributed continuous variables. Data are expressed as frequencies or mean ± standard deviation.
<sup>b</sup>Other: American Indian, Asian, Multiracial, White
\*p < .05
<sup>1</sup>Significant difference from CTL
<sup>2</sup>Significant difference from CUD+
<sup>3</sup>Significant difference from CUD-

### MRI Acquisition

MRI scans were obtained using a Siemens 3.0 Tesla Skyra (Siemens Healthcare, Erlangen, Germany) with a 32-channel head coil. Diffusion-weighted images were acquired with opposite phase encoding along the left-right axis, monopolar diffusion encoding with 128 gradient directions (64 in each phase encoding direction) and 13 b=0 scans (1.8 mm isotropic resolution; repetition time 3650 ms; echo time 87 ms; flip angle 80°; bandwidth 1485 Hz/pixel; single shell maximum *b* value 1500 s/mm^2^, multiband factor = 3, no in-plane acceleration). T1-weighted (T1w) structural scans were acquired using an MPRAGE sequence (0.8 mm isotropic resolution; repetition time 2400 ms; echo time 2.07 ms; inversion time 1000 ms; flip angle 8°; bandwidth 240 Hz/pixel). T2-weighted (T2w) structural scans were acquired using a SPACE sequence (0.8 mm isotropic resolution; repetition time 3200 ms; echo time 565 ms; bandwidth 680 Hz/pixel).

### Structural Data Preprocessing

T1w structural scans were rigidly aligned with each subject’s preprocessed b0 scan using FSL’s FLIRT for preprocessing with the Freesurfer version 6.0 recon-all pipeline (*59*), which includes steps for motion correction, non-uniform intensity normalization, skull stripping, and automated segmentation and parcellation based on the Desikan-Killiany atlas (*60*). The PFC target regions of interest were constructed from parcellations of the superior frontal gyrus (SFG); rostral and caudal middle frontal gyrus (MFG); inferior frontal gyrus (IFG), which contained the pars opercularis, pars orbitalis, and pars triangularis; medial and lateral orbitofrontal cortex (OFC); frontal pole (FP); and rostral and caudal anterior cingulate cortex (ACC).

### Segmentation of Subcortical Seed Regions and Inclusion Mask

Segmentation of the Hb was performed semi-automatically by exploiting the structure’s high myelin content relative to surrounding tissue as described previously (*61*) (Figure 1A). Briefly, each subject’s T2w image was co-registered to the T1w image, and T1w-over-T2w images were generated to obtain myelin contrast maps (*62*). Two individual voxels centered in the left and right Hb region were manually specified, and boundaries were determined automatically by a region growing algorithm with partial volume estimation in the subject’s T1w and T2w images and myelin map. Following visual inspection to ensure anatomical validity, the segmented Hb volumes were shape-optimized and downsampled to individuals’ diffusion MRI space similar to previously described methods for functional MRI analysis (*33*).

To enhance the anatomical accuracy of the estimated Hb streamlines and filter out streamlines originating in adjacent structures including the mediodorsal thalamus, an intermediate inclusion mask was designated in the most anterior aspect of the SM. Selection of the inclusion region was performed in the sagittal plane and guided by the T1w-over-T2w image (Figure 1A, right). A 2 × 2 × 2 voxel region in the resolution of the diffusion space was manually drawn in both hemispheres immediately posterior to the anterior columns of the fornix along the third ventricle.

The area around the inclusion mask contains a handful other fiber pathways, including the anterior thalamic radiations and superolateral medial forebrain bundle. In order to evaluate the specificity of the Hb tractography results, we therefore generated two control seed regions for these respective tracts: the anterior thalamus (AT), which was extracted using Freesurfer’s anteroventral and ventral anterior thalamic nuclei segmentations (*63*) (Figure 1B); and the ventral tegmental area (VTA), which was generated using a previously established atlas of midbrain structures in MNI space (*64*), thresholded at 0.6 to improve specificity, and coregistered to each subject’s b0 scan (Figure 1C).

### Diffusion Data Preprocessing

Diffusion-weighted images were denoised and preprocessed using the *dwipreproc* command in MRtrix3 (*65*), which incorporates FSL tools to correct for eddy current, motion, and susceptibility-induced distortions (*66, 67*). Other corrections were made for B1 field inhomogeneity, and intensity normalization was performed (*68*). Preprocessed diffusion-weighted images were subsequently upsampled to 1.25 mm isotropic voxel resolution. Estimation of the white matter fiber orientation distribution (FOD) function was conducted using the single shell, multi-tissue constrained spherical deconvolution model (*69, 70*). For microstructure analysis of DTI measures, the diffusion tensor was fitted locally in each voxel to generate whole-brain maps of the fractional anisotropy (FA, an index reflecting the coherence of diffusion), mean diffusivity (MD, the bulk averaged/total magnitude of diffusion in all measured directions), axial diffusivity (AD, the magnitude of diffusion in the primary direction), and radial diffusivity (RD, the averaged magnitude of diffusion in the secondary and tertiary orthogonal directions) using the FSL function *dtifit* (*71*). Note that MD may be linked to neurite dispersion and the intracellular space, AD is associated with axon caliber and fiber density, and RD has been previously shown to correlate with histological markers of demyelination (*72*).

### Tractography

Probabilistic tractography was carried out in MRtrix3 using the Second-order Integration over Fiber Orientation Distributions (iFOD2) algorithm (*73*) in the upsampled individual subject space. Each subject’s FOD map was seeded at random within the left and right Hb volumes (and the control regions: AT and VTA) with a step size of 0.9 mm until 100 streamlines (hereby referred to as a “tract”) terminating at the grey/white matter boundary of the PFC were selected per hemisphere. Since, to the best of our knowledge, there are no previous reports indicating the existence of direct contralateral input to the Hb (*27*), the contralateral hemisphere was used as an exclusion region. Parameters for streamline selection were an FOD amplitude threshold of 0.15; 400 mm maximum length; and 20° maximum angle between successive steps. The Anatomically Constrained Tractography framework was included to improve the biological validity of streamline estimates using anatomical priors derived from the 5-tissue type segmented T1-weighted image (*74*), with the additional restriction to the SM inclusion mask for Hb streamlines. Individual tracts were visually inspected to ensure all endpoints were located exclusively in the Hb and PFC, and streamlines that were irrelevant to the intended analysis (e.g., extending posterior or inferior to the Hb through the fasciculus retroflexus) were removed. Tracts from the Hb were segmented into subsections S1, S2, and S3 to assess potential spatial differences in the tissue microstructure in pre-specified anatomical sections. Boundaries were designated in the coronal plane as follows: for S1, immediately anterior to the Hb extending to the anterior SM inclusion mask as described above; for S2, immediately anterior to the genu of the internal capsule extending to the most anterior slice containing the caudate (encompassing the ALIC); and for S3, immediately anterior to the caudate extending to the PFC grey/white matter boundary (Figure 2A). For all tracts, DTI measures were extracted from each voxel along every streamline, and streamline means were averaged across all voxels and between both hemispheres to produce one value per tract for each subject.

For the control regions, streamlines seeded from the VTA were restricted to pass through the ALIC, or S2, as defined above, with exclusion regions in the AT and in the brainstem immediately inferior to the VTA to eliminate overlap with adjacent structures. Streamlines seeded from the AT were restricted to pass through the same ALIC inclusion region.

Visualization of streamlines in the region posterior to and within the ALIC was rendered for topological characterization (Figure 2B). Streamline terminal distributions in the PFC were compared between tracts seeded in each of the subcortical regions of interest (Figure 3A).

To quantify the spatial distinction between Hb and control streamlines in the ALIC, tracts in this region were binarized in each subject’s T1w space. Spatial overlap between the three tracts was computed for each of the three subcortical seed regions as the percentage of the total mask volume contained within each of the other masks, averaged between left and right hemispheres (e.g., the percentage of voxels within in the Hb tract mask that also contain streamlines from the AT) (Figure 2C).

Finally, although tractography was performed with subcortical seed regions and the PFC as the target region, considering that anatomical connectivity with the Hb is unidirectional from the PFC, and tractography is indifferent to directionality, we refer to these projections as “PFC-Hb tracts” to uphold consistency with the previously established anatomy.

### Statistical Analysis

Group comparisons for demographic and drug use measures were performed using chi-square tests for categorical variables, while continuous variables were analyzed using Student’s *t* tests or non-parametric Mann-Whitney U for normal and non-normal distributions, respectively. A comparison of streamline distributions (percent of streamline endpoints) was performed using a 3 (Group: CTL, CUD+, CUD-) × 3 (Seed: Hb, AT, VTA) × 4 (Target: SFG, MFG, IFG, OFC) × 2 (Side: left, right) ANOVA. Group differences in white matter microstructure in each of the three subcortical seed regions were analyzed for each of the four DTI measures with 3 (Group: CTL, CUD+, CUD-) × 2 (Side: left, right) ANOVAs. For these DTI analyses, Bonferroni correction for multiple comparisons was used with an alpha level of .05/12 = .004 to account for four (FA, MD, AD, RD) separate analyses in the three tracts. Additionally, these same analyses were performed separately in S2 to assess specificity in the region of the ALIC where functionally segmented white matter pathways converge yet still maintain topological distinction; we then used as the dependent measure any DTI metrics that survived our multiple comparisons-corrected significance testing in a 3 (Group: CTL, CUD+, CUD-) × 2 (Side: left, right) ANOVA. The same statistical test was performed separately for the other two subsections (S1 and S3) to assess the spatial distribution of any significant DTI effects. For all ANOVAs, significant main effects were followed up with Tukey HSD post hoc tests and simple effects analyses were performed to follow any significant interaction effects. To control for potential covariates, the effects of any demographic, psychometric and non-cocaine use variables that differed between the groups were inspected for their associations with the DTI metrics exhibiting significant group differences (see Table 1). We also assessed correlations between our dependent DTI metrics (that showed significant effects in the analyses above) with drug use and addiction severity variables. Here, for the continuous cocaine use variables (age of first use, duration of regular use, duration of heaviest use, duration of longest abstinence, duration of current abstinence, use in the past 30 days, withdrawal, craving, and severity of dependence), an exploratory factor analysis was conducted to reduce the dimensionality of the data and extract the relevant features in our sample. Factors accounting for a significant amount of the total variance [i.e., sum of squared (SS) loading weights greater than 1] were retained for regression analysis. All statistical tests were performed using R version 3.6.1 (*75*).

## ACKNOWLEDGMENTS

We appreciate the thoughtful discussions with our colleagues, Drs. Patrick Hof, Mark Baxter, Paul Kenny, and Peter Rudebeck, in the Department of Neuroscience at the Icahn school of Medicine at Mount Sinai. We are equally grateful to Dr. Suzanne Haber, in the Department of Psychiatry at Harvard Medical School, for in-depth neuroanatomical discussions.

## Author contributions

S.G.K., N.A-K., J.X., and R.Z.G. conceptualized and designed the research. J.K. and J.X. advised on the methods. S.G.K. performed the data analysis. S.G.K., P-O.G., J.K., J.X., and R.Z.G. interpreted the results. P.M. collected and managed the data. S.G.K. wrote the initial draft of the manuscript. P-O.G., N.A-K., J.K., J.X., and R.Z.G. revised the manuscript.

## Data and materials availability

All data needed to evaluate the conclusions in the paper are present in the paper and/or the Supplementary Materials.

