## Supplementary Figure 1 for "Microstructural Impairments in a Topologically Distinct Prefrontal-Habenular Connection in Cocaine Addiction"

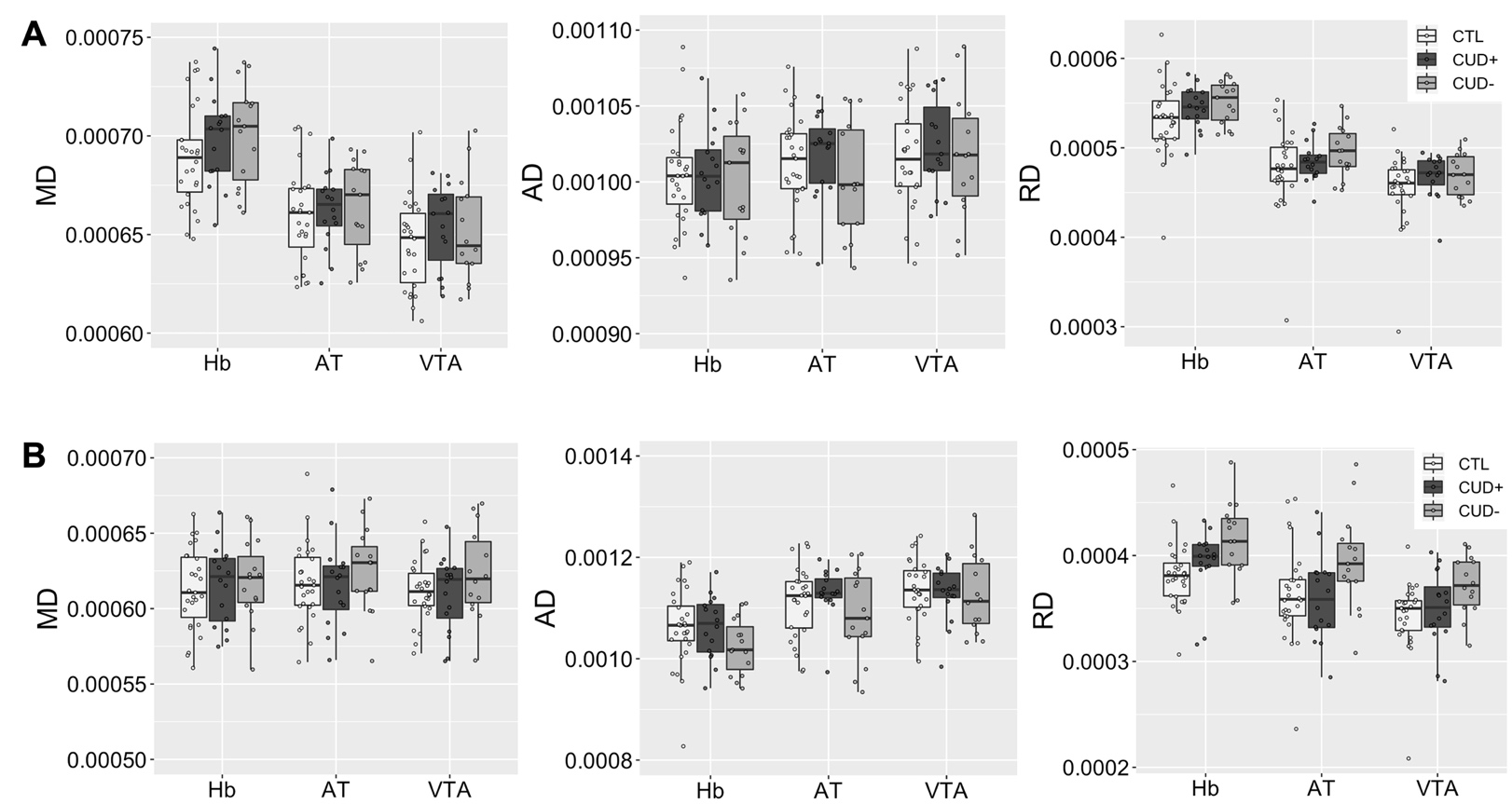

Supplementary Figure 1. Group comparisons of MD, AD, and RD averaged across the entire PFC tract (A) and within the ALIC subsection (B). *p < .05, **p < .01, ***p < .001. MD = mean diffusivity; AD = axial diffusivity; RD = radial diffusivity; Hb = habenula; AT = anterior thalamus; VTA = ventral tegmental area; CTL = control individuals; CUD+ = currently using individuals; CUD- = short-term abstinent individuals. Boxplots represent the median (center line), interquartile range (box), and 1.5x the interquartile range (whiskers).
